# Survival of *Streptococcus anginosus* in blood is linked to hydrogen peroxide production

**DOI:** 10.64898/2026.09.04.748836

**Authors:** Simon Jooß, Verena Vogel, Richard Bauer, Hannah Renner, Stefanie Mauerer, Franziska Rothemund, Eva Gabriel, Parham Sendi, Barbara Spellerberg

**Affiliations:** Institute of Medical Microbiology and Hygiene, University Hospital Ulm, Ulm, Germany; Institute for Infectious Diseases, University of Bern, Bern, Switzerland

## Abstract

*Streptococcus anginosus* is an opportunistic pathogen causing bacteremia, respiratory tract infections, abscesses, odontogenic infections and has recently been associated with gastric cancer. While numerous genes encoding putative virulence factors have been detected in whole genome sequences, a detailed molecular characterization is missing for most of them.

To address putative virulence determinants, we investigated hydrogen peroxide production by *S. anginosus* and assessed its functional significance for survival in human blood. In *Streptococcus pneumoniae* the pyruvate oxidase, encoded by the *spxB* gene, is responsible for H_2_O_2_ production and represents a major virulence factor. Interestingly, *S. anginosus* harbors two genetic variants of the *spxB* gene, one with high homology to the *S. pneumoniae spxB* (sequence identity: approx. 94 %) gene, while the other appears to be specific for *S. anginosus*. Strains carrying the *S. pneumoniae* variant demonstrate higher H_2_O_2_ production (10-20 µM) compared to strains harboring the *anginosus* variant (<1 µM). Incubation of *S. anginosus* strains in human blood showed that bacterial survival was correlated to the amount of hydrogen peroxide production. Assessing H_2_O_2_ production in an S*. anginosus* strain carrying a deletion of the CcpA regulator revealed the control of *spxB* and H_2_O_2_ production by carbon catabolite repression.

In conclusion, we found evidence for the presence of a major *S. pneumoniae* virulence factor in *S. anginosus*, and we were able to demonstrate that it increases the ability of S*. anginosus* to survive in human blood.

## Introduction

*Streptococcus anginosus* is part of the *Streptococcus anginosus* group (SAG), which consists of three bacterial species: *Streptococcus anginosus*, *Streptococcus constellatus* and *Streptococcus intermedius* (Whiley et al. 1999; Grinwis et al. 2010; Lal et al. 2011; Jensen et al. 2013). Bacteria belonging to the SAG are part of the normal human microbiota and were for a long time primarily considered as commensals of mucosal membranes of the oral cavity, the gastrointestinal and the urogenital tract (Poole and Wilson 1976; Asam and Spellerberg 2014; Whiley et al. 1992). However, reports highlighting the pathogenic potential of SAG species have increased in the last decades, with several publications demonstrating their pathogenic role in severe invasive infections (Pilarczyk-Zurek et al. 2022; Qian et al. 2024; Siegman-Igra et al. 2012). These publications clarify the role of SAG as a classical pathobiont, acting as an opportunistic pathogen that can cause a broad spectrum of invasive and polymicrobial infections (Claridge et al. 2001; Asam and Spellerberg 2014; Jiang et al. 2020). Typical infections are abscesses in various organs, including the brain, liver and lungs, as well as chronic pulmonary infections in patients with cystic fibrosis (Parkins et al. 2008; Pilarczyk-Zurek et al. 2022). More recently the ability of *S. anginosus* to colonize even the gastric mucosa, which requires a profound acid resistance has been reported (Masood et al., 2016; Kuryłek et al., 2022). Within this environment, *S. anginosus* contributes to inflammatory responses that can lead to the development of gastric cancer (Fu et al. 2024).

Despite this growing clinical importance of *S. anginosus* our knowledge of virulence factors of this species remains limited, with many having been identified only at the genetic level without experimental validation (Asam and Spellerberg 2014; Kuryłek et al. 2022; Pilarczyk-Zurek et al. 2022). Among these is the pyruvate oxidase, which is responsible for hydrogen peroxide production in streptococcal species (Spellerberg et al. 1996; Ramos-Montañez et al. 2008; Redanz et al. 2018). In *S. pneumoniae*, the pyruvate oxidase is encoded by the gene *spxB*, and intensively studied as an important virulence factor (Spellerberg et al. 1996; Ramos-Montañez et al. 2008, 2010). Pyruvate, which is generated during glycolysis, is converted by SpxB to acetyl phosphate and hydrogen peroxide under aerobic conditions (Ramos-Montañez et al. 2008, 2010). Acetyl-phosphate is utilized in the production of ATP, while acetyl-CoA is essential for cell growth (Ramos-Montañez et al. 2008, 2010). The hydrogen peroxide production by streptococci appears to play a paradoxical role in bacterial competition within the same niche, while also contributing to self-protection against oxidative stress and enhancing bacterial survival (Pericone et al. 2003; Lisher et al. 2017; Redanz et al. 2018). Moreover, H_2_O_2_ produced by streptococcal SpxB induces macrophage cell death modulating the host immune response (Okahashi et al. 2016) and the generated H_2_O_2_ is highly cytotoxic towards brain microglia and other eukaryotic cells (Jennert et al. 2024; Surabhi et al. 2022).

In several streptococcal species the expression of virulence factors is under the control of the carbon catabolite protein A (CcpA) (Stülke and Hillen 1999; Kinkel and McIver 2008; Iyer et al. 2005; Redanz et al. 2020). CcpA functions as a key transcriptional regulator that coordinates carbon catabolism and modulates the expression of genes involved in virulence, stress responses, and metabolic adaptation (Abranches et al. 2008; Roux et al. 2022). It coordinates the balance between growth and damage in pathogenesis (Paluscio et al. 2018). In the presence of elevated glucose levels, CcpA is active and binds to specific DNA sequences known as *cre* sites (catabolite responsive elements), located in the promoter region of the targeted gene, thereby repressing gene expression of these loci (Kietzman and Caparon 2011; Bauer, Mauerer, and Spellerberg 2018). In the context of nutrient-limiting conditions, CcpA is inactive, increasing gene expression of the regulated loci. In *S. anginosus* for example, CcpA controls the expression of hemolysin (Bauer, Mauerer, and Spellerberg 2018). In other streptococci it was demonstrated that CcpA regulates the expression of SpxB (Redanz et al. 2020). Overall, the regulation of *spxB* gene expression by CcpA facilitates bacterial competition for limited carbohydrate sources within a specific host niche by leading to the elimination of bacterial competitors (Jakubovics et al. 2008).

Despite growing interest in *S. anginosus*, many of the potential virulence factors, detected in genetic studies, remain poorly characterized at the functional level, including the pyruvate oxidase. Within this context our study investigates the distribution, regulation and the molecular role of the *S. anginosus* pyruvate oxidase as a potential virulence factor.

## Materials and Methods

### Bacterial cultivation conditions

All bacterial strains, peptides and plasmids used in this study are summarized in Table 1 and Table 2. Streptococcal strains were cultivated on tryptone soy agar plates supplemented with 5% sheep blood (Oxoid) at 37 °C + 5 % CO_2_. Liquid cultivations were carried out in Todd-Hewitt-Yeast (THY) medium (Todd-Hewitt Broth [Oxoid] with 5 % Yeast extract [Gibco]) at 37 °C + 5 % CO_2_. *Escherichia coli* strains were incubated aerobically in lysogeny broth medium (LB-Miller) with 400 µg/mL erythromycin at 37 °C while shaking (160 rpm). *S. anginosus* isolates carrying pAT18-cre-rec_tufA_ were cultivated with 1 µg/ml erythromycin (Serva). *S. anginosus* isolates harboring a spectinomycin resistance were grown in the presence of 120 µg/ml spectinomycin (Merck).

**Table 1:** Wild-type and mutant strains used in this study.

| Strain | Source |
| --- | --- |
| *Streptococcus anginosus* SK52 | ATCC 12395 |
| *Streptococcus anginosus* BSU-Mutant<br>921 (SK52ΔccpA) | Bauer et al., 2018b |
| *Streptococcus anginosus* BSU-Mutant<br>991 (SK52ΔspxB) | This study |
| *Streptococcus anginosus* BSU 1211 | Clinical Isolate of University Hospital Ulm |
| *Streptococcus anginosus* BSU 1212 | Clinical Isolate of University Hospital Ulm |
| *Streptococcus anginosus* BSU 1324 | Clinical Isolate of University Hospital Ulm |
| *Streptococcus anginosus* BSU 1331 | Clinical Isolate of University Hospital Ulm |
| *Streptococcus anginosus* BSU 1338 | Clinical Isolate of University Hospital Ulm |
| *Streptococcus anginosus* BSU 1339 | Clinical Isolate of University Hospital Ulm |
| *Streptococcus anginosus* BSU-Mutant<br>1310 (1339ΔspxB) | This study |
| *Streptococcus anginosus* BSU-Mutant<br>1314 (1339ΔccpA) | This study |
| *Streptococcus intermedius* BSU 1340 | Clinical Isolate of University Hospital Ulm |
| *Streptococcus anginosus* BSU 1356 | Clinical Isolate of University Hospital Ulm |
| *Streptococcus anginosus* BSU 1358 | Clinical Isolate of University Hospital Ulm |
| *Streptococcus anginosus* BSU 1364 | Clinical Isolate of University Hospital Ulm |
| *Streptococcus anginosus* BSU 1366 | Clinical Isolate of University Hospital Ulm |
| *Streptococcus anginosus* BSU 1381 | Clinical Isolate of University Hospital Ulm |
| *Streptococcus anginosus* BSU 1389 | Clinical Isolate of University Hospital Ulm |
| *Streptococcus anginosus* BSU 1401 | Clinical Isolate of University Hospital Ulm |
| *Streptococcus anginosus* BSU 1701 | Clinical Isolate of University of Bern |
| *Streptococcus anginosus* BSU 1712 | Clinical Isolate of University of Bern |
| *Streptococcus anginosus* BSU 1725 | Clinical Isolate of University of Bern |
| *Streptococcus anginosus* BSU 1766 | Clinical Isolate of University of Bern |
| *Streptococcus pneumoniae* BSU 1818 | TIGR4, ATCC BAA-343 |
| *Escherichia coli* 101 | Law et al., 1995 |

**Table 2:**
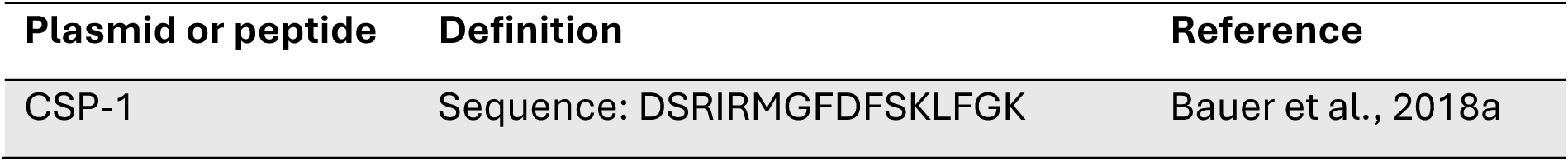
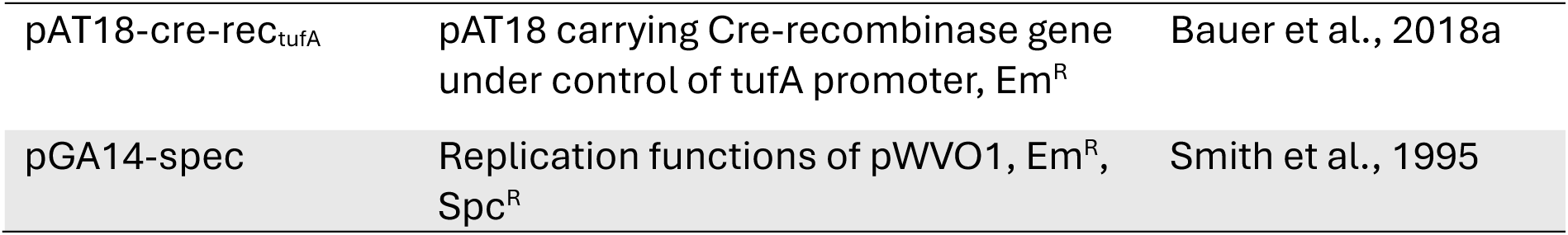
Plasmids and peptides used in this study.

### General DNA-techniques

Genomic DNA was isolated using the QIAamp® DNA Mini Kit (Qiagen) and the GenElute™ Bacterial Genomic DNA Kit (Sigma-Aldrich), while plasmid DNA was extracted using the QIAprep® Spin Miniprep Kit (Qiagen), all according to the manufacturers’ instructions. Polymerase chain reaction (PCR) was performed according to standard protocols for *Taq* polymerase (Roche). PCR cycling conditions consisted of an initial denaturation at 94 °C for 3 min, followed by 33 cycles of denaturation at 94 °C for 30 s, annealing at 50 °C for 30 s, and elongation at 72 °C for 1 – 3 min, with a final elongation step at 72 °C for 7 min. As an alternative to *Taq* polymerase from Roche, PCR was also performed using the AllTaq PCR Master Mix (Qiagen). PCR for fragments larger than 3 kb, was carried out using the Long-expand-system (Expand Long Template PCR System, Roche). Amplified PCR products were cleaned up with NucleoSpin® Gel and PCR Clean-up Kit (Macherey-Nagel). All primers used for this investigation are listed in Supplementary Table S1. Nucleotide sequencing as well as whole genome sequencing were performed by Microsynth Seqlab laboratories (Göttingen, Germany).

### *spxB* screen

To assess the distribution of the *spxB_ang_* and the *spxB_pneu_* gene variants, a PCR-based screening was performed. Primers for the *spxB_pneu_* were designed based on the *S. pneumoniae* TIGR4 *spxB* sequence, whereas primers for the *spxB_ang_* were designed based on the *spxB* sequence of *S. anginosus* SK52.

Primers 1 and 2 amplified the *spxB_pneu_*, while primer pairs 3/4 and 5/6 were specific for the *spxB_ang_*. PCR products were purified and sequenced.

### Hydrogen peroxide measurement

Bacterial hydrogen peroxide production was determined by a commercial kit (Amplex Red Hydrogen Peroxide / Peroxidase Assay Kit, Thermo Fisher Scientific), according to the manufacturer’s instructions. Bacterial cultures were grown for 4 h in THY medium, diluted to an O.D._600nm_ of 1 in 1 ml, washed twice with 1x reaction buffer, and incubated in 1x reaction buffer at 37 °C for 1 h. Supernatants were collected by centrifugation (8800 x g, 2 min, 21 °C), and hydrogen peroxide concentrations were determined by fluorescence measurement (excitation: 542 nm, emission: 592 nm) using a plate reader (Tecan Infinite M Plex). Supernatant of *S. pneumoniae* TIGR4 was used as a positive control, while supernatant obtained from *S. intermedius* served as a negative control.

### Creating ‘markerless’ deletion mutant

Markerless deletion mutants of *S. anginosus* strains were generated as described by Bauer et al. (Bauer, Mauerer, Grempels, et al. 2018), by overlap extension PCR and CSP-mediated transformation.

First, upstream and downstream regions flanking the *ccpA* gene were amplified by *Taq*-PCR with primers containing *loxCC* and *lox71* sequences. A second PCR fragment containing the spectinomycin resistance cassette (spec) was generated from plasmid pGA14-Spc. The resulting fragments were fused by overlap-extension PCR and the full-length construct was subsequently amplified by flanking primers. The resulting linear DNA construct was introduced into bacterial cells via CSP-mediated transformation. For transformation, 100 µL FBS, 10 ng/mL CSP-1, and an overnight culture grown in THY were mixed and incubated at 37 °C for 35 min. Subsequently, 20 µL of the obtained DNA fragment was added, followed by incubation at 37 °C for another 85 min. Cells were then pelleted (8800 x g, 2 min, 21 °C) and plated on THY agar plates supplemented with spectinomycin and incubated for 48 h at 37 °C + 5 % CO_2_. After selection, plasmid pAT18-cre-rec_tufA_ carrying the *Cre-recombinase* gene was introduced via CSP-mediated transformation. Cre-recombinase excised the spectinomycin resistance cassette, leaving a single lox72 site and resulting in a markerless deletion. To cure the plasmid pAT18-cre-rec_tufA_, clones were grown in THY medium for 4 h at 37 °C + 5 % CO_2_ without antibiotic pressure. Clones sensitive to erythromycin were selected and the correct construction of deletions was confirmed by PCR and DNA sequencing.

### Co-culture

To determine whether hydrogen peroxide production leads to a growth advantage in mixed cultures, the hydrogen peroxide producing *S. anginosus* isolate BSU 1339 was inoculated into a co-culture together with the *S. anginosus* type strain SK52. In addition, co-cultures of strain SK52 with *S. anginosus* BSU 1339Δ*spxB* and of SK52 with *S. anginosus* BSU 1339Δ*ccpA* were carried out. All strains were inoculated freshly in THY with an O.D. _600 nm_ of 0.01 and grown for 4 h. Subsequently all strains were adjusted to an O.D. _600 nm_ of 0.1 and the respective mixtures were inoculated in artificial saliva medium (ASM) to mimic physiological conditions. ASM (0.25% mucin, 1% proteose peptone, 1% trypticase peptone, 1% yeast extract, 0.25% KCl, 0.25% glucose) was prepared as previously described (Lin et al. 2025). Samples were drawn after 0, 20, 24 and 28 h, serially diluted and three 10 µl spots per dilution were plated on sheep blood agar plates. After overnight incubation CFU per ml were determined.

### Whole Blood killing assay

To investigate bacterial survival in human blood, a whole blood killing assay was conducted. Overnight cultures were inoculated to an O.D._600nm_ of 0.2 and grown for 90 min. Based on previously obtained growth curves, 10^6^ cells were mixed with 1 ml of human whole blood from healthy volunteers (ethics approval 187-25 of the ethics committee at the University of Ulm). After 0, 2, 4, and 6 h of incubation at 37 °C and shaking at 600 rpm, serially diluted samples were plated on THY agar plates. Following overnight incubation, colony-forming units (CFU) per ml were determined.

### Bioinformatic and statistical analysis

GenBank database was the source for nucleotide sequences (http://www.ncbi.nlm.nih.gov/). Further genetic analysis was performed with CLC Genomics Workbench V22 (https://digitalinsights.qiagen.com/) and Clone Manager V12 (www.scied.com). The putative protein structure of SpxB was predicted based on *spxB* sequence alignments using AlphaFold (Google DeepMind) (Mirdita et al. 2022) and subsequently visualized with iCn3d (National Center for Biotechnology Information, https://www.ncbi.nlm.nih.gov/Structure/icn3d/). GraphPad Prism V10.5.0 was used to visualize data and for statistical analysis (www.graphpad.com).

## Results

### Identification of *spxB* gene variants in *S. anginosus*

Since previous genetic studies in *S. anginosus* (Olson et al. 2013) showed the presence of a pyruvate oxidase gene, we decided to further investigate it as a putative virulence determinant by hydrogen peroxide determinations. A genetic screen identified two distinct *spxB* gene variants among our *S. anginosus* strain collection. One of the variants showed high sequence similarity (93.1 % nucleotide sequence identity, 98.3 % amino acid identity) to the SpxB of *S. pneumoniae* and was designated *spxB_pneu_*, whereas the second variant was labelled as *spxB_ang_*.

Screening of 163 different *S. anginosus* strains showed that the majority of isolates carry a *spxB* gene (79.1 %), and only in a small number of strains no *spxB* gene could be detected (20.9 %) (Table 3). Among strains carrying *spxB*, *spxB_ang_* is the predominant genotype (73.6 %), whereas *spxB_pneu_* is detected less frequently (26.4 %).

**Table 3:** Prevalence of spxB and distribution of *spx*B variants among the investigated *S. anginosus* isolates.

| Characteristic | No. of isolates<br>(Frequency) | Variant | No. of isolates<br>(Frequency) |
| --- | --- | --- | --- |
| Total isolates | 163 |  |  |
| No *spxB* gene | 34 (20.9 %) |  |  |
| *spxB* gene | 129 (79.1 %) | *spxB<sub>ang</sub>* | 95 (73.6 %) |
|  |  | *spxB<sub>pneu</sub>* | 34 (26.4 %) |

The *spxB_pneu_*, identified in *S. anginosus,* shared 93.1 % nucleotide sequence identity with the *spxB* gene of *S. pneumoniae*, whereas both variants shared only approximately 51.9 % sequence identity with each other. At the protein level, SpxB*_ang_* comprises 601 amino acids, compared to 591 amino acids for SpxB*_pneu_*, representing an approximately 10-amino-acid difference in protein length. Despite this difference, the two proteins shared 40.1 % amino acid sequence identity, whereas SpxB*_pneu_* shared 98.3 % amino acid sequence identity with the SpxB pyruvate oxidase of *S. pneumoniae* TIGR4.

Further analysis looked at the up and downstream regions of the two distinct *spxB* gene variants. While the upstream region seems very similar from gene annotation, sequence alignment revealed nucleotide sequence similarity ranging between 72-79 % (Supplementary Table 2, Supplementary Figure 1). The downstream region of the two *spxB* variants differs to a bigger extent, with sequence similarities ranging between 30-40 % (Supplementary Table 3, Supplementary Figure 1). Interestingly, *spxB_ang_* is followed by a gene encoding a lactate oxidase, an enzyme that is also able to produce hydrogen peroxide.

### Protein structure

Since the deduced amino acid sequences revealed considerable heterogeneity between the two variants, we analyzed how this affected the predicted protein structures by AlphaFold. The amino acid sequences of the pyruvate oxidase from *S. anginosus* SK52 harboring the characteristic *spxB_ang_* and *S. anginosus* BSU 1339 carrying the *spxB_pneu_*, were selected as representative. Interestingly, AlphaFold analysis found large parts of the protein structure to be conserved between both variants, despite the sequence heterogeneity (Figure 1). However, the comparison also revealed a predicted structural difference in the region spanning amino acids 549 - 562. In this region, the SpxB*_ang_* variant forms a β-sheet, whereas the SpxB*_pneu_* adopts a coil (Figure 1).

**Figure 1:**
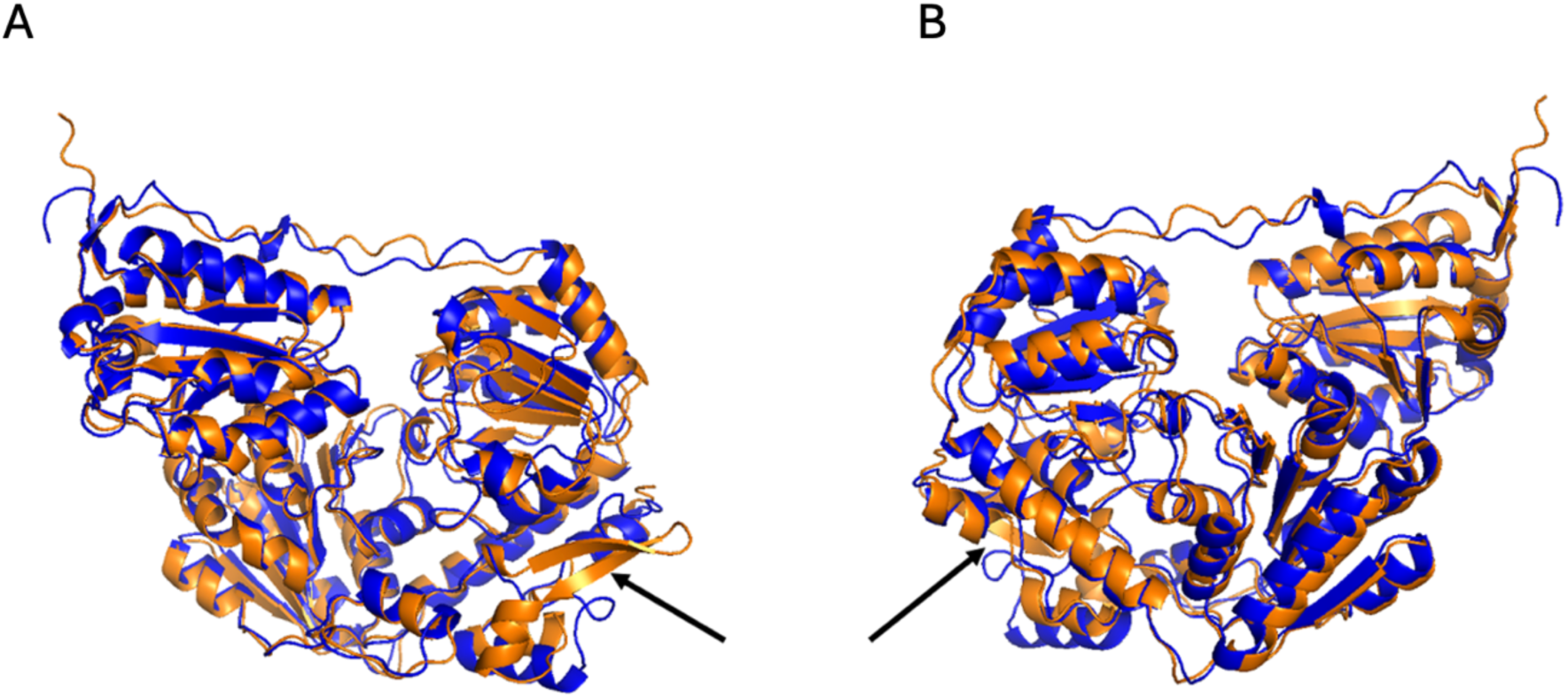
Structural alignment of the predicted SpxB variants from *Streptococcus anginosus.* **A)** Structural alignment of predicted SpxB*_ang_* (orange) and SpxB*_pneu_* (blue) protein structures (AlphaFold) from the front. **B)** Structural alignment of predicted SpxB*_ang_* (orange) and SpxB*_pneu_* (blue) protein structures (AlphaFold) from the back. The arrows indicate the C-terminal region (amino acids 549 – 562), where the biggest predicted structural differences between the two variants are observed. Image was created using iCn3d (https://www.ncbi.nlm.nih.gov/Structure/icn3d/) and the embedded alignment function.

### Hydrogen peroxide production of SpxB*_ang_* and SpxB*_pneu_*

To determine whether the identified *spxB* variants differ at a functional level, hydrogen peroxide production was quantified in 19 chosen *S. anginosus* strains harboring either *spxB_ang_* or *spxB_pneu_*. A significant difference in hydrogen peroxide production between the two variants was detected (Figure 2A). Strains carrying *spxB_pneu_* produce significantly higher levels of H_2_O_2_ than those carrying *spxB_ang_*.

**Figure 2:**
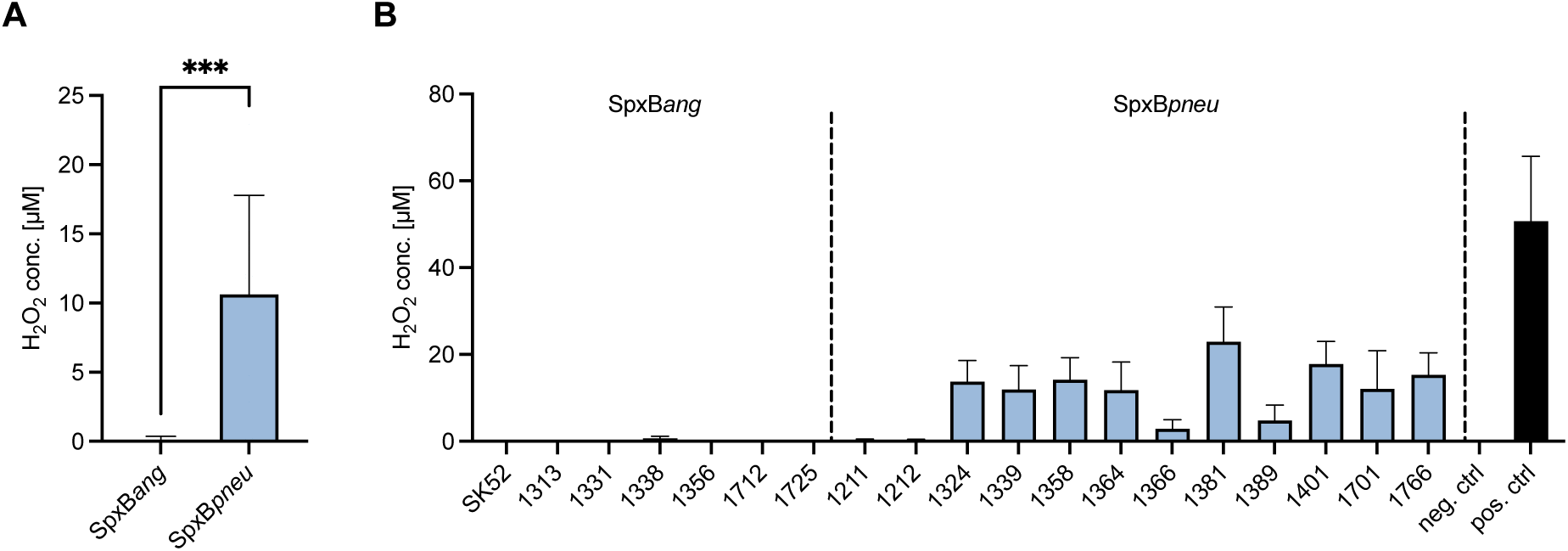
Hydrogen peroxide production by *Streptococcus anginosus* strains. *S. anginosus* strains were grown in THY broth for 4 h and subsequently incubated in assay buffer for 1 h before hydrogen peroxide concentration was determined using the Amplex Red assay. **A)** Mean hydrogen peroxide production of all isolates carrying either *spxB_ang_* (n=7) or *spxB_pneu_* (n=12). Statistical significance was assessed using the Mann-Whitney U test (\*\*\**p* < 0.001). **B)** Hydrogen peroxide production of all investigated *S. anginosus* strains, grouped according to their respective SpxB variant. Data are illustrated as mean + SD from at least five independent experiments.

Strains carrying *spxB_ang_* generally exhibited minimal or no hydrogen peroxide production, whereas the majority of strains carrying *spxB_pneu_* produced markedly higher levels of hydrogen peroxide (Figure 1B). Among these strains, *S. anginosus* BSU 1381, generated the highest level of hydrogen peroxide with up to 24 µM. However, not all of the strains carrying *spxB_pneu_* demonstrated a pronounced H_2_O_2_ production, BSU 1211 and BSU 1212 had hydrogen peroxide production levels comparable to strains carrying *spxB_ang_*.

### Influence of SpxB and CcpA on hydrogen peroxide production

To confirm the importance of SpxB for hydrogen peroxide production, deletion mutants of *spxB_pneu_* and *spxB_ang_* were created in the strains BSU 1339 and SK52 respectively. As expected, strain BSU 1339 produced higher levels of hydrogen peroxide than strain SK52, and the deletion of *spxB* abolished the hydrogen peroxide production in both strains, regardless of the respective gene variant (Figure 3). These results confirm that the pyruvate oxidase SpxB is responsible for hydrogen peroxide production in *S. anginosus* under aerobic conditions.

**Figure 3.**
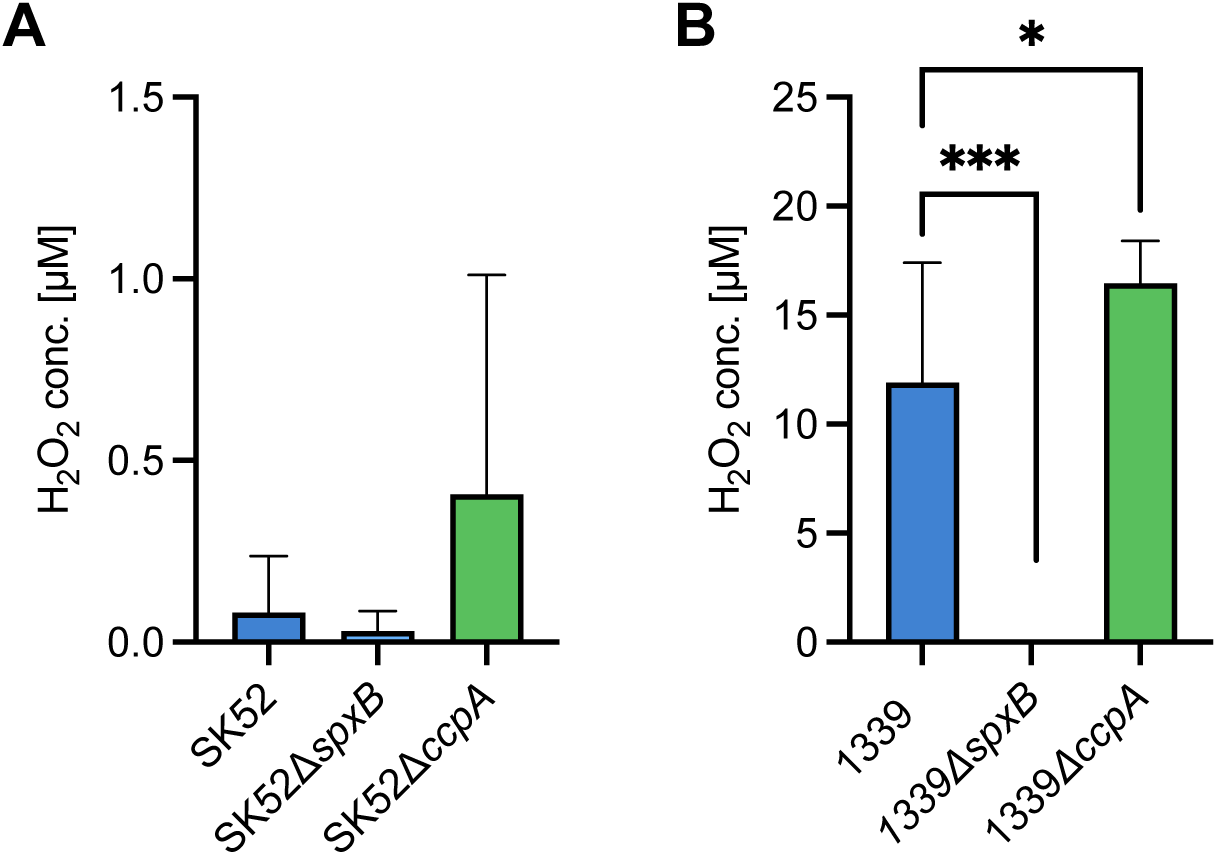
Hydrogen peroxide production of *Streptococcus anginosus* wild-type strains and their corresponding isogenic Δ*spxB* and Δ*ccpA* deletion mutants. **A)** Hydrogen peroxide production of *S. anginosus* SK52 and its corresponding isogenic deletion mutants. **B)** Hydrogen peroxide production of *S. anginosus* BSU 1339 and its corresponding isogenic deletion mutants. Data are presented as mean + SD from at least five independent experiments. Statistical significance was assessed using Mann-Whitney U test (\**p* < 0.05, \*\**p* < 0.01, \*\*\**p* < 0.001).

To investigate whether regulatory mechanisms contribute to the differences in hydrogen peroxide production the influence of carbon catabolite repression was examined. The carbon catabolite regulator CcpA has been shown to regulate hemolysin expression in *S. anginosus* and SpxB in other streptococcal species (Bauer, Mauerer, and Spellerberg 2018; Redanz et al. 2020).

To determine to what extent the observed hydrogen peroxide production is regulated by CcpA, *ccpA* deletion mutants of *S. anginosus* SK52, carrying *spxB_ang_*, and *S. anginosus* BSU 1339, carrying the *spxB_pneu_* were analyzed (Figure 3). Deletion of *ccpA* resulted in an increased hydrogen peroxide production in *S. anginosus* SK52, compared with the corresponding wild-type strain. However, this difference was not statistically significant, presumably due to the high standard deviations observed for H₂O₂ levels, which remained comparatively low relative to the hydrogen peroxide levels produced by BSU 1339. In the background of strain BSU 1339, H_2_O_2_ was significantly increased in the Δ*ccpA* deletion mutant indicating that CcpA acts as a negative regulator of SpxB in *S. anginosus*.

### Analysis of putative *cre* sites

Given that a *ccpA* deletion increased hydrogen peroxide production, we investigated putative CcpA binding sites in the *spxB* promotor region. In all strains, analyzed for H_2_O_2_ production (Figure 1B), the region upstream of the *spxB* gene was analyzed for putative *cre* motifs, that are recognized by CcpA using the FIMO tool of the Meme Suite (https://meme-suite.org/meme/doc/fimo.html). This analysis revealed three distinct 16-bp sequences consistent with the consensus motif of a *cre* binding site (Figure 4A). While all strains harbouring *spxB_ang_* have an identical putative *cre* binding site, two distinct *cre* binding sites were identified upstream of *spxB_pneu_*. Interestingly, the different motifs were very clearly associated with different hydrogen peroxide production levels (Figure 4B).

**Figure 4:**
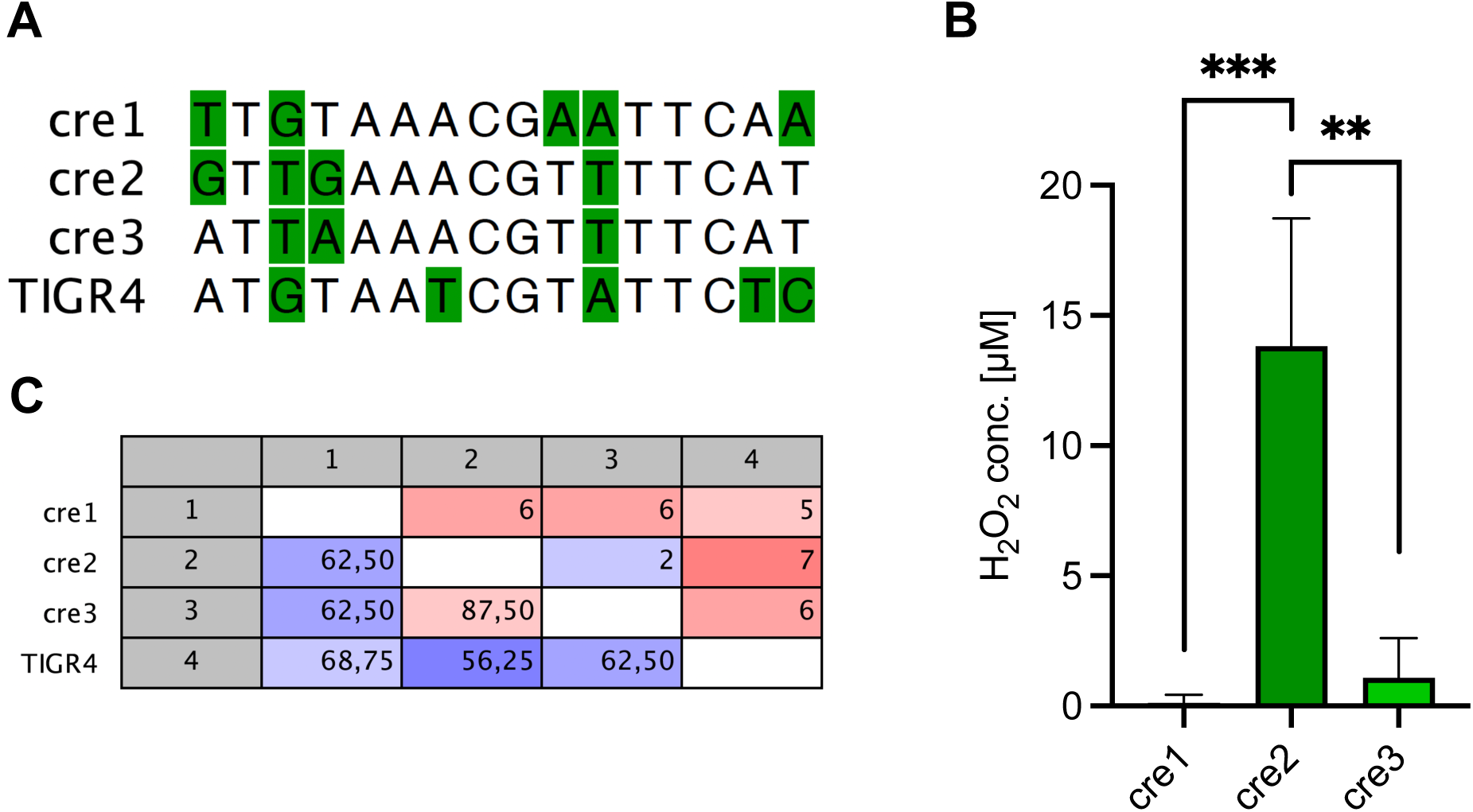
Carbon catabolite repression regulates hydrogen peroxide production. **A)** Sequence alignment of the predicted carbon catabolite control protein A (CcpA) binding sites upstream of *spxB* in representative *S. anginosus* isolates and the corresponding *cre* site of *S. pneumoniae* TIGR4. Illustration was created using with CLC Genomics Workbench V22 (https://digitalinsights.qiagen.com/). **B)** Mean hydrogen peroxide production of strains carrying identical predicted CcpA binding sites. Data are presented as mean + SD from at least five independent experiments. Statistical significance was assessed using Mann-Whitney U test (\**p* < 0.05, \*\**p* < 0.01, \*\*\**p* < 0.001). **C)** Pairwise comparison of the predicted *cre* site sequences, showing the number of nucleotide differences (upper triangle) and the corresponding pairwise sequence identities (lower triangle). Analysis and visualization were performed with CLC Genomics Workbench V22 (https://digitalinsights.qiagen.com/).

All seven strains harboring the *S. anginosus spxB* gene variant exhibited the identical putative *cre* binding site TTGTAAACGAATTCAA (*cre*1), which was associated with low or absent H_2_O_2_ production.

Some *S. anginosus* strains (BSU 1211 and 1212) carrying *spxB_pneu_* showed only a minimal H_2_O_2_ production with concentrations below 0.5 µM H_2_O_2_. For both of these strains, the putative *cre* site ATTAAAACGTTTTCAT (*cre*3) was found. This putative *cre* site is also found in the isolate BSU 1366, which has also a very limited H_2_O_2_ production with 2.855 µM. All other isolates carrying *spxB_pneu_* showed higher levels of H_2_O_2_ (4.75 - 24.304 µM) and harbored the different putative *cre* site GTTGAAACGTTTTCAT (*cre*2). While for *cre3* the mean production is at 1.07 µM H_2_O_2_, the mean production of isolates carrying *cre2* is significantly higher with 15.49 µM. Among the analyzed strains carrying *spxB_pneu_*, 75 % were high producers and possessed the GTTGAAACGTTTTCAT *cre2* site, while 25 % possessed the ATTAAAACGTTTTCAT *cre3* site.

The alignment of all putative *cre* sites identified upstream of *spxB* revealed a highly conserved core region at nucleotide positions 5 – 11, whereas the terminal regions exhibited greater sequence variability (Figure 4A). Pairwise sequence comparison demonstrated the highest sequence identity between the two putative *cre* sites upstream of *spxB_pneu_* (87.5 %) (Figure 4C). In contrast, the putative *cre* site associated with the *spxB_ang_* shared higher sequence identity with the low H_2_O_2_-producing *spxB_pneu_* (68.8 %) than with the high H_2_O_2_-producing *spxB_pneu_* (62.5 %). The reported *S. pneumoniae* TIGR4 *cre* site exhibited moderate sequence identity with all putative *cre* sites identified in *S. anginosus*, sharing the highest sequence identity with the *spxB_ang_* cre site (68.8 %).

In addition to the sequence variation, the position of the putative *cre* sites relative to the *spxB* start codon differed between the identified variants. The putative *cre* site of the low producing *spxB_ang_* variant (*cre*1) was located 94 bp upstream of the *spxB* start codon, whereas the putative *cre* site of the high producing *spxB_pneu_* variant (*cre*2) was located 70 bp upstream. In addition, the low producing *spxB_pneu_ cre* site (*cre*3) was located 74 bp upstream of the start codon. By comparison, the reported *S. pneumoniae* TIGR4 *cre* site is located 49 bp upstream of the *spxB* start codon.

Taken together, strains carrying putative *cre* sites located further upstream of the *spxB* start codon exhibited lower hydrogen peroxide production.

### Bacterial competition in co-culture

Hydrogen peroxide contributes to bacterial competition and self-protection in streptococci (Pericone et al., 2003; Lisher et al., 2017; Redanz et al., 2018). To investigate its role in bacterial competition under conditions mimicking the oral cavity, the low hydrogen peroxide producer, *S. anginosus* SK52, carrying *spxB_ang_*, was co-incubated in ASM with *S. anginosus* BSU 1339, carrying *spxB_pneu_*. In addition, co-incubations were performed with the corresponding *S. anginosus* BSU 1339Δ*spxB* and *S. anginosus* BSU 1339Δ*ccpa* mutants (Figure 5).

**Figure 5:**
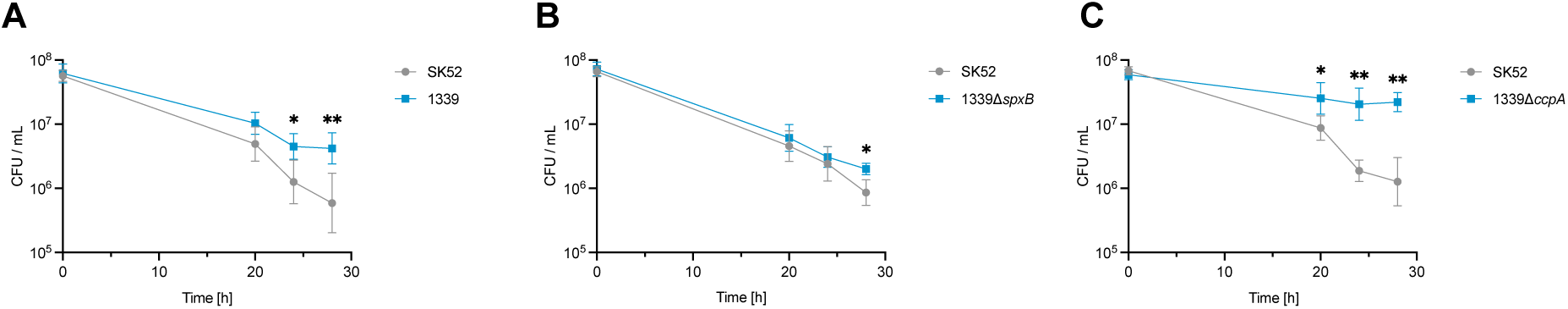
Influence of hydrogen peroxide production on intraspecies competition. *S. anginosus* SK52 was co-cultured with *S. anginosus* BSU 1339 **A)** wild-type, **B)** the isogenic Δ*spxB* deletion mutant, or **C)** the isogenic Δ*ccpA* deletion mutant in artificial saliva medium (ASM). Samples were collected after 20, 24, and 28 h, and bacterial survival was determined by enumeration of colony-forming units (CFU). Data are presented as mean ± SD from at least five independent experiments. Statistical significance was assessed using Mann-Whitney U test (\**p* < 0.05, \*\**p* < 0.01, \*\*\**p* < 0.001).

Viability of *S. anginosus* SK52 remained largely comparable across all conditions, whereas the viable counts of *S. anginosus* BSU 1339 and its mutants varied, according to the level of H_2_O_2_ production.

During co-incubation of both wild-type strains, *S. anginosus* BSU 1339, which produces higher levels of H_2_O_2_ than *S. anginosus* SK52, exhibited increased viability compared to *S. anginosus* SK52 (Figure 5A). In contrast, the viable counts of H_2_O_2_-deficient mutant *S. anginosus* BSU 1339Δ*spxB* were comparable to that of S*. anginosus* SK52 (Figure 5B). The high H_2_O_2_ producing mutant *S. anginosus* BSU 1339Δ*ccpA* exhibited significantly increased viable counts, reaching the highest cell densities among all the tested strains (Figure 5C).

These results indicate that increased hydrogen peroxide production was associated with enhanced growth of the producing strain in co-culture.

### Whole Blood Killing Assay

To assess the potential role of the pyruvate oxidase as a virulence factor, a whole blood killing assay was performed in human blood (Figure 6). Survival rates of *S. anginosus* SK52 and *S. anginosus* BSU 1339 were compared with their isogenic *spxB* and *ccpA* deletion mutants (Figure 6) after 2, 4 and 6 hours of incubation in human blood. Deletion of *ccpA* significantly increased the survival of both strains in comparison to the wild types, consistent with the increased hydrogen peroxide production of the *ccpA* deletion mutants.

**Figure 6:**
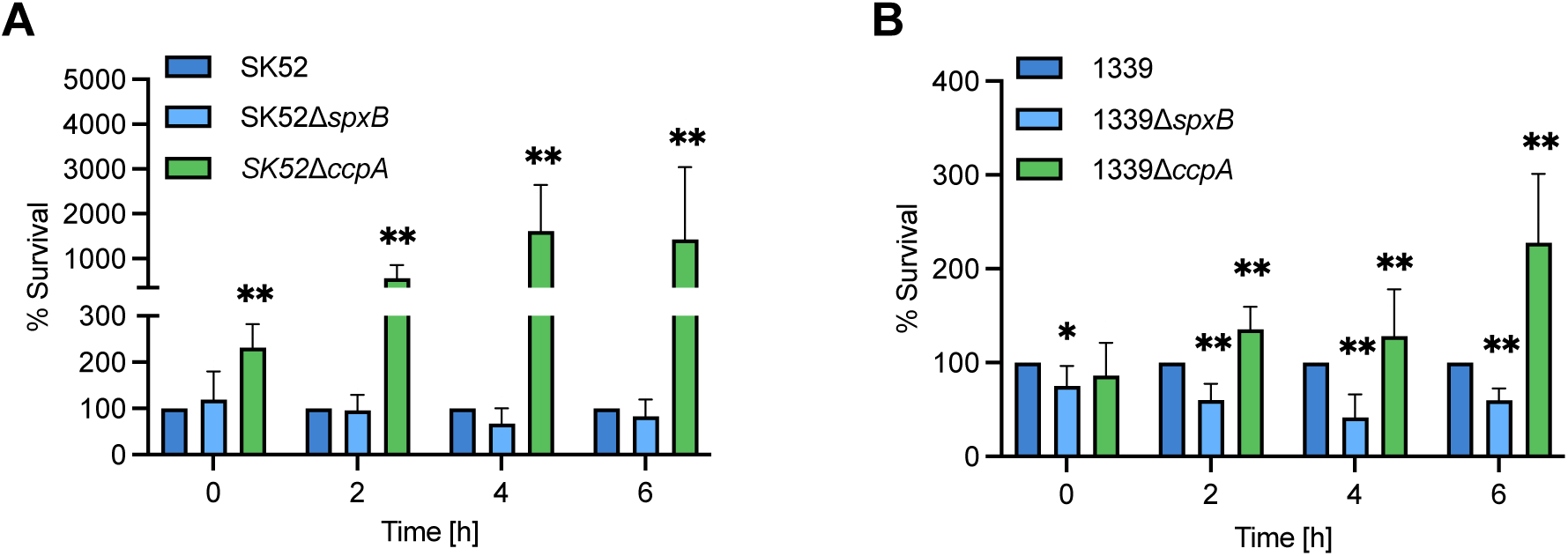
Survival of *S. anginosus* in human whole blood. **A)** Survival of *S. anginosus* SK52 and **B)** BSU 1339 and their corresponding isogenic Δ*spxB* and Δ*ccpA* deletion mutants in human whole blood. Samples were collected every 2 h over a 6 h incubation period, and bacterial survival was determined by enumeration of colony-forming unit (CFU). CFU values of the deletion mutants were normalized to those of their respective wild-type strain. Data are presented as mean + SD from at least five independent experiments. Statistical significance was assessed using Mann-Whitney U test (\**p* < 0.05, \*\**p* < 0.01, \*\*\**p* < 0.001).

The importance of hydrogen peroxide production for survival in blood could be further substantiated by deletion of *spxB*, which significantly decreased the survival of *S. anginosus* BSU 1339 (Figure 6B). As expected, no significant effect was observed for *S. anginosus* SK52Δ*spxB*, likely reflecting the already low baseline hydrogen peroxide production of this strain (Figure 6A). Consistent with these findings, the increase in survival following *ccpA* deletion was more pronounced in *S. anginosus* SK52Δ*ccpA*, than in S*. anginosus* BSU 1339Δ*ccpA*.

In summary higher hydrogen peroxide production was associated with substantially increased survival in human blood supporting a potential role for SpxB as a virulence factor of *S. anginosus*.

## Discussion

In many pathogenic streptococcal species, virulence factors are well characterized, allowing a deeper understanding about the molecular basis for the development of serious infections. This is, however, not the case in *S. anginosus,* where our knowledge about virulence factors is currently very limited. Through genetic screens in over 150 clinical *S. anginosus* strains, we detected two distinct *spxB* gene variants, encoding pyruvate oxidases, that are responsible for markedly different levels of hydrogen peroxide production. One of these variants displays a very close genetic similarity to the *S. pneumoniae* pyruvate oxidase, which is a major pneumococcal virulence factor (Ramos-Montañez et al. 2008; Spellerberg et al. 1996). In our investigation we characterized the role of the different *S. anginosus* pyruvate oxidases for survival of bacteria in blood and mixed bacterial cultures and we investigated how hydrogen peroxide production is regulated through the carbon catabolite regulator CcpA.

### SpxB variants and H_2_O_2_ production

Amino acid alignments of the two *S. anginosus spxB* gene variants, show that the *spxB_pneu_* variant shares 98 % homology with the *spxB* gene present in *S. pneumoniae*, while the deduced protein sequence of the *spxB_ang_* variant shares only about 40 % homology to *S. pneumoniae.* This genetic difference is matched by highly different levels of hydrogen peroxide production. Strains carrying SpxB*_ang_* generally produced H_2_O_2_ close to the limit of detection of our assay at about 0.5 µM or below, in contrast to strains harboring SpxB*_pneu_* with H_2_O_2_ production levels of up to 25 µM. Since *S. pneumoniae* is known to produce exceedingly high levels of hydrogen peroxide (Pericone et al., 2003), this close sequence similarity may very well account for the high H_2_O_2_ production of strains harboring SpxB*_pneu_.* However, in comparison to *S. pneumoniae* and other oral streptococcal species hydrogen peroxide production in *S. anginosus* still appears to be modest. *S. pneumoniae*, for example, is known to produce up to 1 mM H₂O₂ after 30 minutes of aerobic incubation, while *S. gordonii* produces approximately 180 µM after 1 hour and *S. sanguinis* produces lower levels, at approximately 120 µM after 1 hour of aerobic incubation (Pericone et al., 2003; Xu et al., 2014). These moderate production levels raised the question, if the two SpxB variants are both linked to hydrogen peroxide production. Since the deletion of the respective *spxB* genes resulted in a complete abolishment of hydrogen peroxide production for each variant, our results clearly confirm that both *spxB* variants are responsible for hydrogen peroxide production in *S. anginosus* under aerobic conditions. To assess how the strikingly different protein sequences of both variants that are however associated with the conserved function of H_2_O_2_ production via the oxidation of pyruvate may affect protein structure, an alpha fold analysis was conducted. Surprisingly, despite the observed sequence heterogeneity the structural predictions of both variants looked very similar (Figure 1). The major difference lies in the region encompassing amino acids 549 to 570: while a β-sheet is predicted for SpxB*_ang_*, a coiled structure is predicted for SpxB*_pneu_*. In other bacterial species the structure of the pyruvate oxidase has been elucidated by traditional x-ray crystallography. In *Aerococcus viridans* structure analysis demonstrated SpxB to be organized as a homotetramer consisting of 4 identical subunits (Juan et al. 2007). These four subunits are arranged in a V-shape configuration with the open side facing inward, converging towards the center to form the catalytic center (Juan et al. 2007). Although the structural organization of SpxB in *S. anginosus* has not yet been elucidated, a similar homotetrameric structure appears to be possible according to the AlphaFold predictions. The high structural similarity of the two predicted proteins suggests that differences in hydrogen peroxide production are unlikely to result primarily from major differences in SpxB structure or catalytic architecture. Instead, they may reflect differences in bacterial metabolism, environmental conditions, or transcriptional regulation. Streptococci possess additional hydrogen peroxide-producing enzymes, including lactate oxidase, which may further contribute to isolate-specific production levels (Redanz et al. 2018). Although pyruvate oxidase is considered the principal source of hydrogen peroxide during aerobic exponential growth, lactate oxidase may become more relevant under oxygen-limited conditions and during stationary phase. Consequently, the ecological niche and associated metabolic conditions of each species may influence the relative contribution of these pathways. *S. anginosus* is frequently isolated from oxygen-poor abscesses, whereas *S. pneumoniae* predominantly colonizes the comparatively oxygen-rich respiratory tract. The higher hydrogen peroxide production observed in *S. pneumoniae* may therefore reflect not only differences in transcriptional regulation but also adaptation to the distinct oxygen and nutrient environments encountered by these species (Cheng et al. 2018; Cundell et al. 1995; Parkins et al. 2008; Pilarczyk-Zurek et al. 2022).

### CcpA-mediated regulation of *spxB*

One of the regulators exerting transcriptional control in *S. anginosus* that may affect *spxB* gene expression and contribute to these phenotypic differences is CcpA. It is a key regulator in streptococci, known to control carbohydrate-related pathways, by binding to *cre* sites upstream of its target genes (Carvalho et al. 2011). CcpA-mediated gene regulation has previously been characterized in several streptococci, such as *S. pyogenes*, *S. mutans*, *S. agalactiae*, *S. pneumoniae* and *S. anginosus* (Iyer et al. 2005; Abranches et al. 2008; Shelburne et al. 2008; Moye et al. 2014; Bauer, Mauerer, and Spellerberg 2018; Roux et al. 2022). In *S. anginosus* CcpA regulates transcription of the hemolysin genes and a typical *cre* site (GCGAAAGCGCTTTTTT) was identified upstream of the β-hemolysin gene cluster (Bauer, Mauerer, and Spellerberg 2018). For SpxB, CcpA has also been described as a regulator, in *S. sanguinis, S. gordonii* and *S. pneumoniae*(Hu et al. 2023; Zheng, Itzek, et al. 2011; Zheng, Chen, et al. 2011).

To investigate the effect of CcpA on H_2_O_2_ production in *S. anginosus*, deletion mutants of the *ccpA* gene in *S. anginosus* BSU 1339 (*spxB_pneu_*) and SK52 (*spxB_ang_*) were analyzed. In both *S. anginosus* strains, the mutants lacking *ccpA* exhibited a significantly increased H_2_O_2_ production, compared to the wild-type strain expressing *ccpA* (Figure 5). This increase is most likely attributable to the loss of CcpA-mediated repression of *spxB*, confirming that SpxB is under the control of CcpA in *S. anginosus*, irrespective of the SpxB gene variant.

To further characterize the CcpA-mediated regulation, putative *cre* binding sites upstream of *spxB* were sequenced and analyzed in the 19 strains that were previously selected for functional pyruvate oxidase analysis. This led to the identification of three different *cre* binding sites characteristic for specific levels of hydrogen peroxide production (Figure 4). The putative *cre* site associated with *spxB_ang_* (TTGTAAACGAATTCAA) was associated with low hydrogen peroxide production, whereas two distinct putative *cre* sites were identified upstream of *spxB_pneu_*. Of these, GTTGAAACGTTTTCAT was associated with high hydrogen peroxide production, whereas ATTAAAACGTTTTCAT was associated with comparatively lower production levels (Figure 4C). Further analysis by sequence alignments revealed a conserved core region in the center of all the identified *cre* binding sites, while the flanking sequences show greater variability, which may lead to differences in the binding affinity of CcpA (Figure 4). Comparison of the identified sequences with the *cre* site that is present upstream of the hemolysin genes (GCGAAAGCGCTTTTTT) (Bauer, Mauerer, and Spellerberg 2018) showed similar conserved nucleotide residues. In particular, the length of 16 bases is identical to the identified SpxB *cre* sites and the core region from base 4 to 8 is highly similar, underlining their putative role as CcpA binding sites.

Previous studies have demonstrated that the regulatory effect of CcpA depends not only on the nucleotide sequence, but also on the position of the *cre* site relative to the promoter and transcription start site (Deutscher et al. 2006). *Cre* sites located close to the transcription initiation region may interfere with RNA polymerase binding, thereby repressing transcription.

In summary, our results indicate that the hydrogen peroxide production in *S. anginosus* is controlled by CcpA and we were able to reveal a very specific association between particular *cre* sites and the level of hydrogen peroxide production.

Interestingly, strains carrying the putative *cre* site ATTAAAACGTTTTCAT exhibited hydrogen peroxide production levels comparable to those of strains carrying the putative *cre* site TTGTAAACGAATTCAA. However, despite their overall low hydrogen peroxide production, substantial variation was observed through the strain carrying the putative *cre* site ATTAAAACGTTTTCAT. As these strains share the same *spxB* nucleotide sequence, *cre* site, and *cre* site position, these observations suggest that additional factors may contribute to the observed phenotypic variation. Besides differences in *spxB* regulation, other enzymes involved in hydrogen peroxide metabolism, such as lactate oxidase, may also contribute to the observed phenotype. In streptococci lactate oxidase has been shown to represent an additional source of H_2_O_2_, particularly during stationary growth under low oxygen conditions (Redanz et al. 2018). However, whether lactate oxidase contributes to H_2_O_2_ production in *S. anginosus* remains to be determined.

### Hydrogen peroxide production affects survival rates of *S. anginosus* in mixed cultures and human blood

SpxB has been described to provide a colonization advantage to streptococci (Regev-Yochay et al. 2007; Zhu et al. 2018). Within the mucosal microbiota hydrogen peroxide release into the extracellular environment helps to eliminate other bacterial pathogens and to ensure bacterial survival (J. A. Imlay et al. 1988; Redanz et al. 2018; James A. Imlay 2003; Pericone et al. 2003; Hernandez-Morfa et al. 2023). Within this context we investigated how H_2_O_2_ production affected survival of *S. anginosus* in mixed cultures (Figure 5). These experiments demonstrated that *S. anginosus* survival is clearly linked to H_2_O_2_ production levels. In particular, the CcpA mutant strain of BSU 1339 with the highest H_2_O_2_ levels outcompeted the SK52 wildtype strain significantly. Our data strengthen the putative role of hydrogen peroxide production by *S. anginosus* in conferring a colonization advantage.

To investigate the role of hydrogen peroxide production in invasive infections beyond the colonization stage, we studied the survival of *S. anginosus* in human blood. For these experiments strain SK52, a low-level hydrogen peroxide producer, and the high-H₂O₂-producing strain, BSU 1339 was chosen and compared to their isogenic deletion mutants SK52Δ*spxB*, SK52Δ*ccpA* and BSU 1339Δ*spxB,* BSU 1339Δ*ccpA.* Our results indicate that survival in human blood is associated with the H₂O₂ production levels of the respective strains. Deletion of *spxB* significantly decreased survival in BSU 1339 but had no significant effect on SK52. This difference is consistent with a contribution of hydrogen peroxide production to BSU 1339 survival, but not to survival of the low-level hydrogen peroxide-producing SK52 isolate. In contrast, deletion of *ccpA* resulted in significantly increased survival of both BSU 1339 and SK52 (Figure 6). This effect was especially strong for SK52. Given that CcpA regulates hydrogen peroxide production but also numerous metabolic and virulence-associated pathways in streptococci, additional factors may contribute to the increased survival of the *ccpA* mutants (Bai et al. 2019; Carvalho et al. 2011; Kinkel and McIver 2008).

The improved bacterial survival associated with increased H_2_O_2_ production may result from a modulation of the host immune system. Reduced H_2_O_2_ production has been linked to enhanced phagocytosis by neutrophils, as shown for *S. sanguinis* and *S. pneumoniae* (Bättig and Mühlemann 2008; Syk et al. 2014). Furthermore, the induction of neutrophil death and NET formation by H_2_O_2_ has been reported for *S. sanguinis* (Sumioka et al. 2017). Hydrogen peroxide may also oxidize hemoglobin and promote heme degradation, releasing free iron that can contribute to hydroxyl radical formation and cytotoxicity (Alibayov et al. 2022). In addition, H_2_O_2_ can inhibit inflammasome activation by oxidizing apoptosis-associated speck-like protein and related components, reducing IL-1β and IL-18 release and thereby impairing early immune clearance (Erttmann and Gekara 2019). These mechanisms contribute to enhanced bacterial survival in human blood and may thus play an important part in the development of invasive *S. anginosus* infections. Overall, this data substantiates the role of SpxB as a virulence factor of *S. anginosus*.

Taken together, we identified two *spxB* variants associated with distinct hydrogen peroxide production levels. The *S. anginosus* variant is associated with low H₂O₂ production, whereas strains carrying the *S. pneumoniae* variant produce higher levels of H₂O₂. We were also able to show that *spxB* expression is under carbon catabolite repression via CcpA, and that specific *cre* sites are associated with the amount of hydrogen peroxide produced. Furthermore, we substantiated the role of the pyruvate oxidase as a virulence factor of *S. anginosus* by demonstrating the fitness advantage it provides in mixed cultures and its important role for survival in human blood.

## Authors Contributions

Genetic screens were conducted by FR, SJ, HR, RB and VV. Hydrogen peroxide concentration determination was performed by SJ, HR and VV. Mutants were created by SJ, HR, FR, RB and VV. Whole blood killing experiments were established and performed by SJ, EG, RB. Co-culture experiments were performed by SM. Conceptualization was done by RB, VV and BS. The first draft was written by SJ, VV and BS and reviewed by all authors.

## Acknowledgments

VV thanks the Medical Faculty of Ulm University for funding via the project LSSH1000.36 within the Hertha-Nathorff-Programme.

## Ethics Declaration

Experiments involving human blood were approved within ethics approval 187-25 of the ethics committee at the University of Ulm. All other experiments were performed in accordance with German laws.

## Data Availability Statement

The raw data supporting the conclusions of this article will be made available by the authors, without undue reservation.

