## Supplemental Tables 1, 2 and 3, and supplemenatal Figure 1 for "Survival of *Streptococcus anginosus* in blood is linked to hydrogen peroxide production"

### **Supplementary Table 1: Primers used in this study**

| Number |  | Oligonucleotide / primer | Sequence (5’ – 3’) |
| --- | --- | --- | --- |
| 1 |  | spxB_screen2_fwd | TTATGGCTTTGAAGGTGGAG |
| 2 |  | spxB_screen2_rev | CCTCCCAAGTCTATGAATG |
| 3 |  | spxB_screen_fwd | CTCCTGAATTGGCTTATTATG |
| 4 |  | spxB_screen_rev | CCTTCAGCACTACTAGTTTG |
| 5 |  | spxB_screen3_fwd | GCTGCTGTTGCTATGCTTCG |
| 6 |  | spxB_screen3_rev | TATTCCCTCTACTCCCATTC |
| 7 |  | ccpA_F1_fwd | TCCCAAGGATTTTCAGAGTC |
| 8 |  | ccpA_F2_rev | AATCCGCCATTTGGTACATGTG |
| 9 |  | lox66_spec_rev | TACCGTTCGTATAATGTATGCTATACGAAGTTATATGCCTAGCAGGTCGATTTTCG |
| 10 |  | lox71_spec_fwd | TACCGTTCGTATAGCATACATTATACGAAGTTATTTAAATGGCATTGGTACCC |
| 11 |  | 764_spec_fwd | GTAACCATTCTCCATAAATAAATTC |
| 12 |  | spec_rev | GAATTTATTTATGGAGAATGGTTAC |
| 13 |  | spxB_pneu_seq_rev | CTTGCATCACCGCTGCAAG |
| 14 |  | spxB_pneu2_F1_fwd | GCTTCTGCAGCAATGCTCAAC |
| 15 |  | spxB_pneu2_F2_rev | CCAAGAAGAGGCGGAATGG |
| 16 |  | 62CV_ccpA_F1_fwd | GCTGGCAAGGCTGCTCTG |
| 17 |  | 1218_1339_ccpA_F1_fwd | CGCACTAGCGCGATCTACT |
| 18 |  | 1219_1339_ccpA_F1_rev | GCATACATTATACGAACGGTACTGTATCGTCTGTGTTCAT |
| 19 | | 1220_1339_ccpA_F2_fwd | TATAATGTATGCTATACGAACGGTACAGGCACAGTTGATGTGGAG |
| 20 | | 1221_1339_ccpA_F2_rev | GACATGAGCTAATACCACTTC |
| 21 | | 1250_spxB_Angi_seq_rev | CCAGGACCAGCAGATCCAAG |
| 22 | | 1251_spxBAngi_seq_fwd | GAATGGGAGTAGAGGGAATA |

### **Supplementary Table 2:** Description of the upstream region of the *spxB* gene of H_2_O_2_ measured and whole genome sequenced *S. anginosus* isolates.

| S. anginosus strain | size upstream region | gene annotations upstream | | | | spxB |
| --- | --- | --- | --- | --- | --- | --- |
| SK52 | 6885 bp | helix-turn-helix domain-containing protein | hypothetical protein | DUF6287 domain-containing protein | ldcB, LD-carboxypeptidase LdcB/DacB | *spxB_ang_* |
| BSU 1313 | 5829 bp | helix-turn-helix domain-containing protein |  | DUF6287 domain-containing protein | ldcB, LD-carboxypeptidase LdcB/DacB | *spxB_ang_* |
| BSU 1338 | 5945 bp | helix-turn-helix domain-containing protein |  | DUF6287 domain-containing protein | ldcB, LD-carboxypeptidase LdcB/DacB | *spxB_ang_* |
| BSU 1211 | 6604 bp | helix-turn-helix domain-containing protein | SH3 domain-containing protein | DUF6287 domain-containing protein | ldcB, LD-carboxypeptidase LdcB/DacB | *spxB_pneu_* |
| BSU 1212 | 6658 bp | helix-turn-helix domain-containing protein | SH3 domain-containing protein | DUF6287 domain-containing protein | ldcB, LD-carboxypeptidase LdcB/DacB | *spxB_pneu_* |
| BSU 1324 | 6651 bp | helix-turn-helix domain-containing protein | SH3 domain-containing protein | DUF6287 domain-containing protein | ldcB, LD-carboxypeptidase LdcB/DacB | *spxB_pneu_* |
| BSU 1339 | 6651 bp | helix-turn-helix domain-containing protein | SH3 domain-containing protein | DUF6287 domain-containing protein | ldcB, LD-carboxypeptidase LdcB/DacB | *spxB_pneu_* |
| BSU 1364 | 6614 bp | helix-turn-helix domain-containing protein | SH3 domain-containing protein | DUF6287 domain-containing protein | ldcB, LD-carboxypeptidase LdcB/DacB | *spxB_pneu_* |
| BSU 1366 | 6604 bp | helix-turn-helix domain-containing protein | SH3 domain-containing protein | DUF6287 domain-containing protein | ldcB, LD-carboxypeptidase LdcB/DacB | *spxB_pneu_* |
| BSU 1381 | 6651 bp | helix-turn-helix domain-containing protein | SH3 domain-containing protein | DUF6287 domain-containing protein | ldcB, LD-carboxypeptidase LdcB/DacB | *spxB_pneu_* |
| BSU 1389 | 6614 bp | helix-turn-helix domain-containing protein | SH3 domain-containing protein | DUF6287 domain-containing protein | ldcB, LD-carboxypeptidase LdcB/DacB | *spxB_pneu_* |
| BSU 1401 | 6651 bp | helix-turn-helix domain-containing protein | SH3 domain-containing protein | DUF6287 domain-containing protein | ldcB, LD-carboxypeptidase LdcB/DacB | *spxB_pneu_* |
| BSU 1701 | 6651 bp | helix-turn-helix domain-containing protein | SH3 domain-containing protein | DUF6287 domain-containing protein | ldcB, LD-carboxypeptidase LdcB/DacB | *spxB_pneu_* |

### **Supplementary Table 3:** Description of the downstream region of the *spxB* gene of H_2_O_2_ measured and whole genome sequenced *S. anginosus* isolates.

| S. anginosus strain | spxB | gene annotations downstream | | | | | size downstream region |
| --- | --- | --- | --- | --- | --- | --- | --- |
| SK52 | *spxB_ang_* | lctO, Lactate oxidase; | 6-phospho-beta-glucosidase | response regulator transcription factor | sensor histidine kinase | ABC transporter permease | 5642 bp |
| BSU 1313 | *spxB_ang_* | lctO, Lactate oxidase; | 6-phospho-beta-glucosidase | response regulator transcription factor | sensor histidine kinase | ABC transporter permease | 5642 bp |
| BSU 1338 | *spxB_ang_* | lctO, Lactate oxidase; | 6-phospho-beta-glucosidase | response regulator transcription factor | sensor histidine kinase | ABC transporter permease | 5642 bp |
| BSU 1211 | *spxB_pneu_* | RNA-directed DNA polymerase | RNA-directed DNA polymerase | SemiSWEET family transporter | 6-phospho-beta-glucosidase |  | 5674 bp |
| BSU 1212 | *spxB_pneu_* | hypothetical protein -region downstream seems different | ABC transporter ATP-binding protein | alpha/beta fold hydrolase | energy-coupling factor transporter transmembrane component T |  | 5384 bp |
| BSU 1324 | *spxB_pneu_* | RNA-directed DNA polymerase | RNA-directed DNA polymerase | SemiSWEET family transporter | 6-phospho-beta-glucosidase |  | 5674 bp |
| BSU 1339 | *spxB_pneu_* | RNA-directed DNA polymerase | RNA-directed DNA polymerase | SemiSWEET family transporter | 6-phospho-beta-glucosidase |  | 5675 bp |
| BSU 1364 | *spxB_pneu_* | RNA-directed DNA polymerase | RNA-directed DNA polymerase | SemiSWEET family transporter | 6-phospho-beta-glucosidase |  | 5676 bp |
| BSU 1366 | *spxB_pneu_* | RNA-directed DNA polymerase | RNA-directed DNA polymerase | SemiSWEET family transporter | 6-phospho-beta-glucosidase |  | 5674 bp |
| BSU 1381 | *spxB_pneu_* | RNA-directed DNA polymerase | RNA-directed DNA polymerase | SemiSWEET family transporter | 6-phospho-beta-glucosidase |  | 5682 bp |
| BSU 1389 | *spxB_pneu_* | RNA-directed DNA polymerase | RNA-directed DNA polymerase | SemiSWEET family transporter | 6-phospho-beta-glucosidase |  | 5674 bp |
| BSU 1401 | *spxB_pneu_* | RNA-directed DNA polymerase | RNA-directed DNA polymerase | SemiSWEET family transporter | 6-phospho-beta-glucosidase |  | 5674 bp |
| BSU 1701 | *spxB_pneu_* | RNA-directed DNA polymerase | RNA-directed DNA polymerase | SemiSWEET family transporter | 6-phospho-beta-glucosidase |  | 5675 bp |

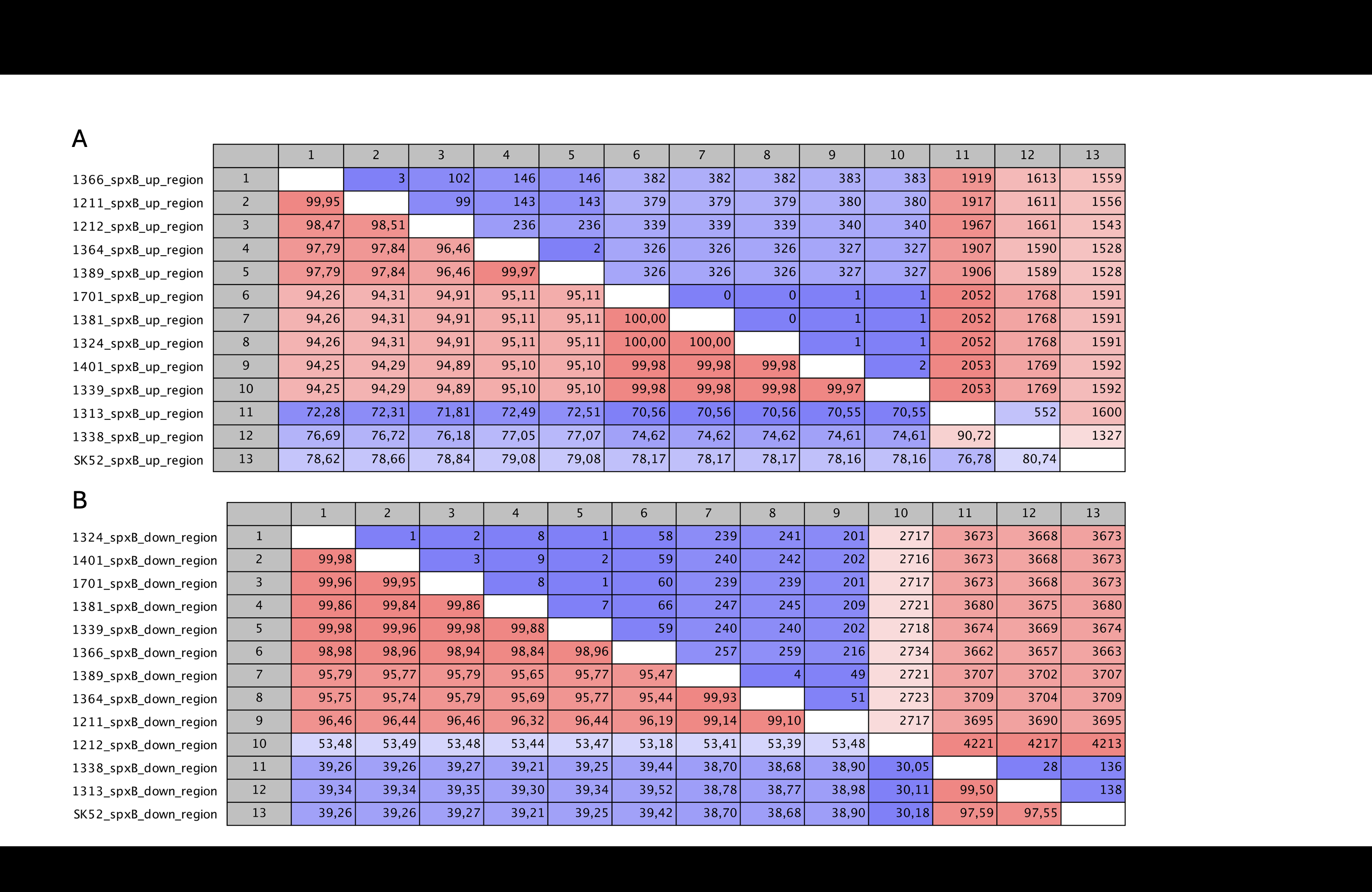

### **Supplementary Figure 1: Pairwise Comparison matrix of the up and downstream region of *spxB*.**

Pairwise comparison of the up (A) and downstream (B)sequence of *spxB* of the 13 whole genome sequenced *S. anginosus* isolates, showing the number of nucleotide differences (upper triangle) and the corresponding pairwise sequence identities (lower triangle). Analysis and visualization were performed with **CLC Genomics Workbench V22 (**<https://digitalinsights.qiagen.com/>**).**
